# DUAL ACTION ELECTROCHEMICAL BANDAGE OPERATED by a PROGRAMMABLE MULTIMODAL WEARABLE POTENTIOSTAT

**DOI:** 10.1101/2024.03.22.586346

**Authors:** Ibrahim Bozyel, Derek Fleming, Won-Jun Kim, Peter F. Rosen, Suzanne Gelston, Dilara Ozdemir, Suat U. Ay, Robin Patel, Haluk Beyenal

## Abstract

Electrochemical bandages (e-bandages) can be applied to biofilm-infected wounds to generate reactive oxygen species, such as hypochlorous acid (HOCl) or hydrogen peroxide (H_2_O_2_). The e-bandage-generated HOCl or H_2_O_2_ kills biofilms *in vitro* and in infected wounds on mice. The HOCl-generating e-bandage is more active against biofilms *in vitro*, although this distinction is less apparent *in vivo*. The H_2_O_2_-generating e-bandage, more than the HOCl-generating e-bandage, is associated with improved healing of infected wounds. A strategy in which H_2_O_2_ and HOCl are generated alternately—for dual action—was explored. The goal was to develop a programmable multimodal wearable potentiostat (PMWP) that could be programmed to generate HOCl or H_2_O_2_, as needed. An ultralow-power microcontroller unit managed operation of the PMWP. The system was operated with a 260-mAh capacity coin battery and weighed 4.6 grams, making it suitable for small animal experiments or human use. The overall cost of a single wearable potentiostat was $6.50 (USD). The device was verified using established electrochemical systems and functioned comparably to a commercial potentiostat. To determine antimicrobial effectiveness, PMWP-controlled e-bandages were tested against clinical isolates of four prevalent chronic wound bacterial pathogens, methicillin-resistant *Staphylococcus aureus* (MRSA), *Pseudomonas aeruginosa, Acinetobacter baumannii*, and *Enterococcus faecium*, and one fungal pathogen of emerging concern, *Candida auris*. PMWP-controlled e-bandages exhibited broad-spectrum activity against biofilms of all study isolates tested when programmed to deliver HOCl followed by H_2_O_2_. These results show that the PMWP operates effectively and is suitable for animal testing.

## INTRODUCTION

Chronic wound healing is a significant health care challenge, afflicting between 5 and 7 million Americans annually and incurring an estimated yearly cost of $50 billion in the United States alone (Copeland and Purvis, 2017). A leading factor contributing to chronic wound persistence is infection by biofilm-producing bacteria. Biofilms are communities of microorganisms protected by a self-secreted matrix known as extracellular polymeric substance (EPS). It has been estimated that more than 90% of chronic wound infections are biofilm-based (Attinger and Wolcott, 2012; Römling and Balsalobre, 2012). Biofilms offer advantages for pathogen survival, including physical protection from the host immune system and antimicrobials, retention of water, tolerance to desiccation, nutrient sorption and storage, high extracellular enzymatic activity, adhesion to the infection site, and coordination of virulence factor expression through quorum sensing (Flemming and Wingender, 2010; Karatan and Watnick, 2009; Rumbaugh et al., 2009). Biofilms complicate chronic wounds by inducing a perpetual state of inflammation, hindering healing, causing collateral damage by the immune system, and protecting pathogens from antimicrobials and host defenses (Watters et al., 2013; Zhao et al., 2013). Additionally, biofilms raise the effective minimum inhibitory concentrations of antimicrobials against pathogens (Bjarnsholt et al., 2013). This is particularly problematic when biofilms are formed by antimicrobial-resistant pathogens. Therefore, it is prudent to explore alternatives to traditional antimicrobial treatments that can broadly combat biofilm infections while circumventing the emergence of antimicrobial-resistant strains.

One method of combatting antimicrobial-resistant pathogens in biofilms is using biocidal molecules. Biocides are commonly included in antiseptics and disinfectants as primary active compounds (McDonnell and Russell, 1999; Ortega-Peña et al., 2017). Sodium hypochlorite (NaOCl), hypochlorous acid (HOCl), hydrogen peroxide (H_2_O_2_), and ozone (O_3_) are biocides that have been highlighted for their antimicrobial efficacy (Haws et al., 2018; Robson et al., 2007a, 2007b). Their clinical implementation in wound infection control has been challenging, as low concentrations are needed to avoid host toxicity, while continuous presence at the infection site is necessary for effective microbe control. As a strategy for overcoming these hurdles, these compounds can be electrochemically generated on wounds at clinically determined time intervals without interruption, enabling wound infection control by eradicating pathogens in wound beds (Jones and Joshi, 2021).

Wound beds naturally contain small quantities of H_2_O_2_ and HOCl (Atiyeh et al., 2009; Robson et al., 2007b; Roy et al., 2006) produced by immune cells to combat pathogens, lessen inflammation, and remove dead tissue. Low levels of H_2_O_2_ can enhance wound healing by promoting keratinocyte and fibroblast migration to wound beds (Zhu et al., 2017). Continuous low-level H_2_O_2_ production is necessary for efficient wound healing applications, as H_2_O_2_ is chemically unstable and decomposes over time (Černŷ et al., 2018). HOCl is another biocide created during the innate immune response (Robson et al., 2007b). Myeloperoxidase, active during inflammation, converts H_2_O_2_ generated by neutrophils into HOCl (Vissers and Winterbourn, 1995). HOCl is a powerful biocidal agent for application in wound infection treatment (Fleming et al., 2023; McMahon et al., 2020; Ortega-Peña et al., 2017).

An electrochemical bandage (e-bandage) is a device that can be applied to a biofilm-infected wound to generate reactive oxygen species, such as HOCl or H_2_O_2_. It consists of three electrodes (working, counter, and reference) immersed in a hydrogel. The working electrode can generate HOCl or H_2_O_2_ based on polarization potential. Our research group showed that HOCl or H_2_O_2_ electrochemically generated by e-bandages can be effectively used to treat biofilms *in vitro* and in infected murine wounds (Kletzer et al., 2023a; Mohamed et al., 2023; Raval et al., 2023; Tibbits et al., 2022). The *in vitro* activity of the HOCl-producing e-bandage was, in general, greater than that of the H_2_O_2_-producing e-bandage *in vitro*, albeit not necessarily *in vivo* (Fleming et al., 2023), while the H_2_O_2_-producing e-bandage augmented wound healing more than the HOCl-producing e-bandage (Raval et al., 2023). Another approach would be to generate H_2_O_2_ and HOCl alternately for dual action. With available technology, however, such dual action can only be tested *in vitro* using commercial potentiostats, as there is no wearable potentiostat suitable for animal (or human) testing of intermittent H_2_O_2_ and HOCl generation by an e-bandage.

Here, the development of a programmable multimodal wearable potentiostat (PMWP) that can be programmed to generate HOCl or H_2_O_2_ selectively in e-bandages via polarization is described. The device is programmable, allowing the duration of HOCl or H_2_O_2_ treatment to be variable. Experiments were designed to 1) test the electronic performance of the PMWP, 2) test the electrochemical performance of PMWP-controlled e-bandages, 3) test the HOCl and H_2_O_2_ generation of PMWP-controlled e-bandages, and 4) demonstrate the activity of PMWP-controlled e-bandage treatment of *in vitro* biofilms formed by clinical isolates of common wound pathogens. The PMWP was designed to operate an e-bandage that can be applied like a standard adhesive wound dressing. Its programmability allows generation of HOCl or H_2_O_2_ for desired times (e.g., for antimicrobial activity and wound healing).

## MATERIALS and METHODS

### e-Bandage

An e-bandage is a three-electrode electrochemical system saturated with hydrogel in which the working electrode (WE) and the counter electrode (CE) are separated by an additional thin layer of hydrogel (**Fig. 1A**). A Ag/AgCl-coated silver wire acting as a pseudo reference electrode (RE) is placed between the WE and CE, and the layers are separated by cotton fabric. **Fig. 1B** shows a schematic of the three-electrode system with readout circuitry. The control amplifier (CA) regulates the voltage level on the CE. The positive terminal of the CA is controlled by the variable voltage channel of a digital-to-analog converter (DAC). A feedback branch is formed between the negative and output terminals of the transimpedance amplifier (TIA) to convert current to voltage for measurement purposes. The TIA feedback branch includes resistors and capacitors for filtering and current-to-voltage conversion processes. Another DAC channel drives the positive terminal of the TIA, allowing the current flow direction between the CE and WE to be adjusted and the cell potential of the potentiostat to be set. Moreover, the voltage level at the positive terminal of the TIA is used for coupling the output of the TIA to fit in the input range of the analog-to-digital converter (ADC).

**Fig. 1.**
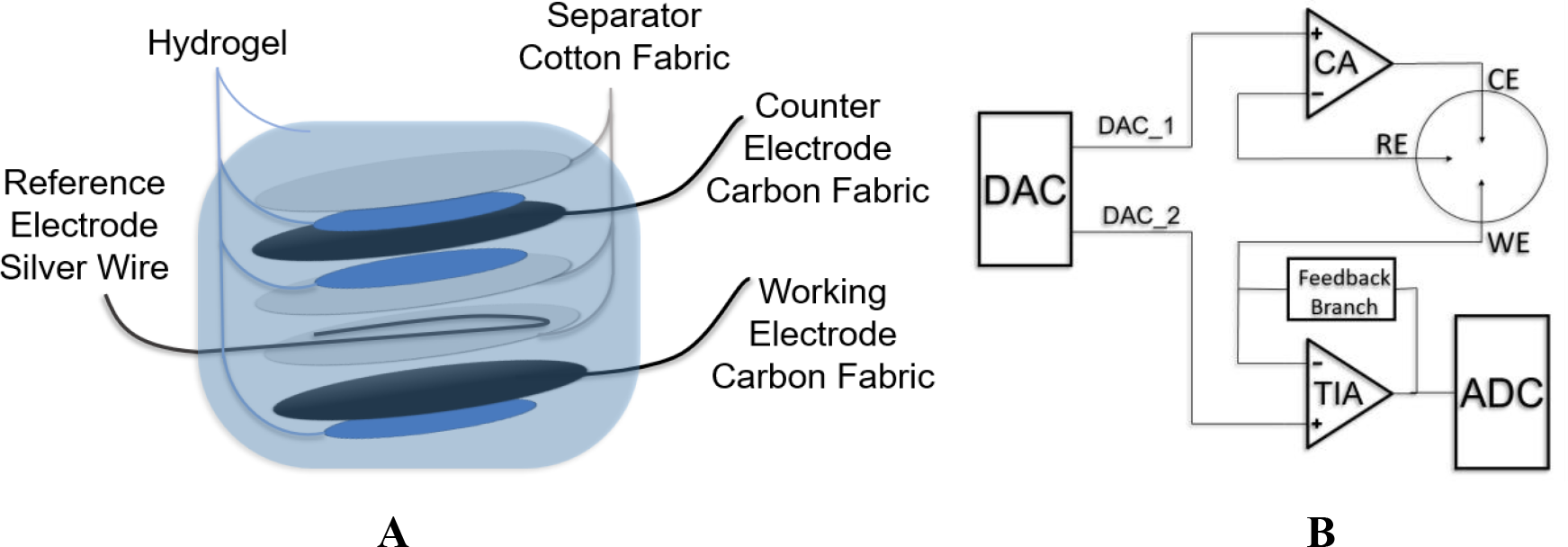
**A)** Schematic of an e-bandage. **B)** Schematic of a 3-electrode electrochemical cell with readout circuitry. DAC, TIA, and ADC refer to the digital-to-analog converter, the transimpedance amplifier, and the analog-to-digital converter, respectively.

The hydrogel made of xanthan gum contains dissolved O_2_ and NaCl. When the WE is polarized to 1.5 V_Ag/AgCl_, chloride ions are oxidized to chlorine, which further reacts with water to generate HOCl (Equation 1).

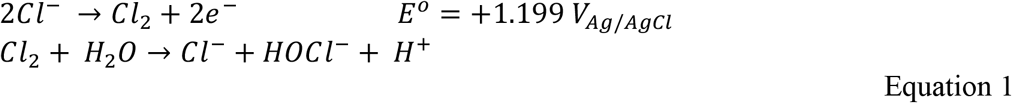

Equation 1 However, when the WE is polarized to -0.6 V_Ag/AgCl_, H_2_O_2_ is produced as a result of partial oxygen reduction (Equation 2) (Raval et al., 2021; Sultana et al., 2015):

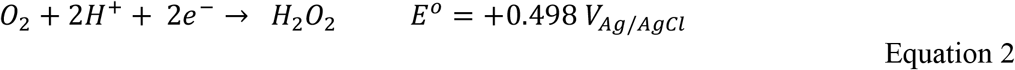

## Programmable Multimodal Wearable Potentiostat

### Power and analog part

The programmable multimodal wearable potentiostat (PMWP) was designed using an integrated circuit with two operational amplifiers (Texas Instruments, LPV542). Each operational amplifier consumes 490 nA of quiescent current and can supply a maximum of 14 mA of output current when the supply voltage is 3.3 V. The LPV542 has a low input bias current of 100 fA and a low input offset voltage of ±1 mV. The PMWP is operated using a 3.0-V coin battery and is designed to operate using supply voltages between 2.0 V and 5.5 V. If the battery voltage drops below 2.0 V, the PMWP detects this and ceases operation. A boost switch mode voltage regulator (Texas Instruments, TPS61023) maintains the system supply voltage at 3.3 V; it consumes 0.9 µA of quiescent current and provides 92% efficiency. The pass-through feature of the TPS61023 allows the PMWP to work even if the battery is overcharged.

### Microcontroller

The controller unit of the PMWP is an ultralow-power series 32-bit ARM Cortex^®^ M4 core CPU with a floating point unit (FPU) (STM32L431). This microcontroller unit (MCU) has several power modes. Generally, power modes of MCUs are categorized as “sleep,” “standby,” and “shutdown.” The STM32L series has three additional stop modes that enable precise control of specific peripherals. The STOP1 mode was selected to run chronoamperometry. In this mode, two DAC channels and a GPIO block remain active. An on-chip 12-bit ADC (0.805-mV resolution on 3.3-V reference voltage) samples TIA output. The transimpedance amplifier feedback branch includes a 100-Ω resistor. The non-inverting input of the TIA is fed by the DAC channel-1 to set the WE voltage through feedback. Consequently, the current generated within the electrochemical cell is amplified 100-fold, enabling detection of microampere-level changes. A 1-kΩ resistor star network is used as a dummy cell, and each resistor pin is connected to the WE, RE and CE positions for life cycle tests (Devices, 2018).

### Programming and production

Electronic design software (KiCAD) was used to design the schematic and the printed circuit board (PCB) layout of the PMWP. Two-layered PCBs were produced by OSH-Park. Integrated circuits (ICs), an ON/OFF switch, connectors and passives were placed on the top layer while a battery holder was placed on the bottom layer of the PCB (**Fig. 2**). Components were manually soldered using a hot plate. Software validation and programming were completed using STMCubeMX and STMCubeIDE tools. Pinout, configurations, and a clock were assigned through a STMCubeMX user interface. Embedded C code was written in STMCubeIDE. **Fig. 2** shows a photograph of a PMWP with a connected e-bandage. The entire schematic of the PMWP design is shown in **Fig. S1**.

**Fig. 2.**
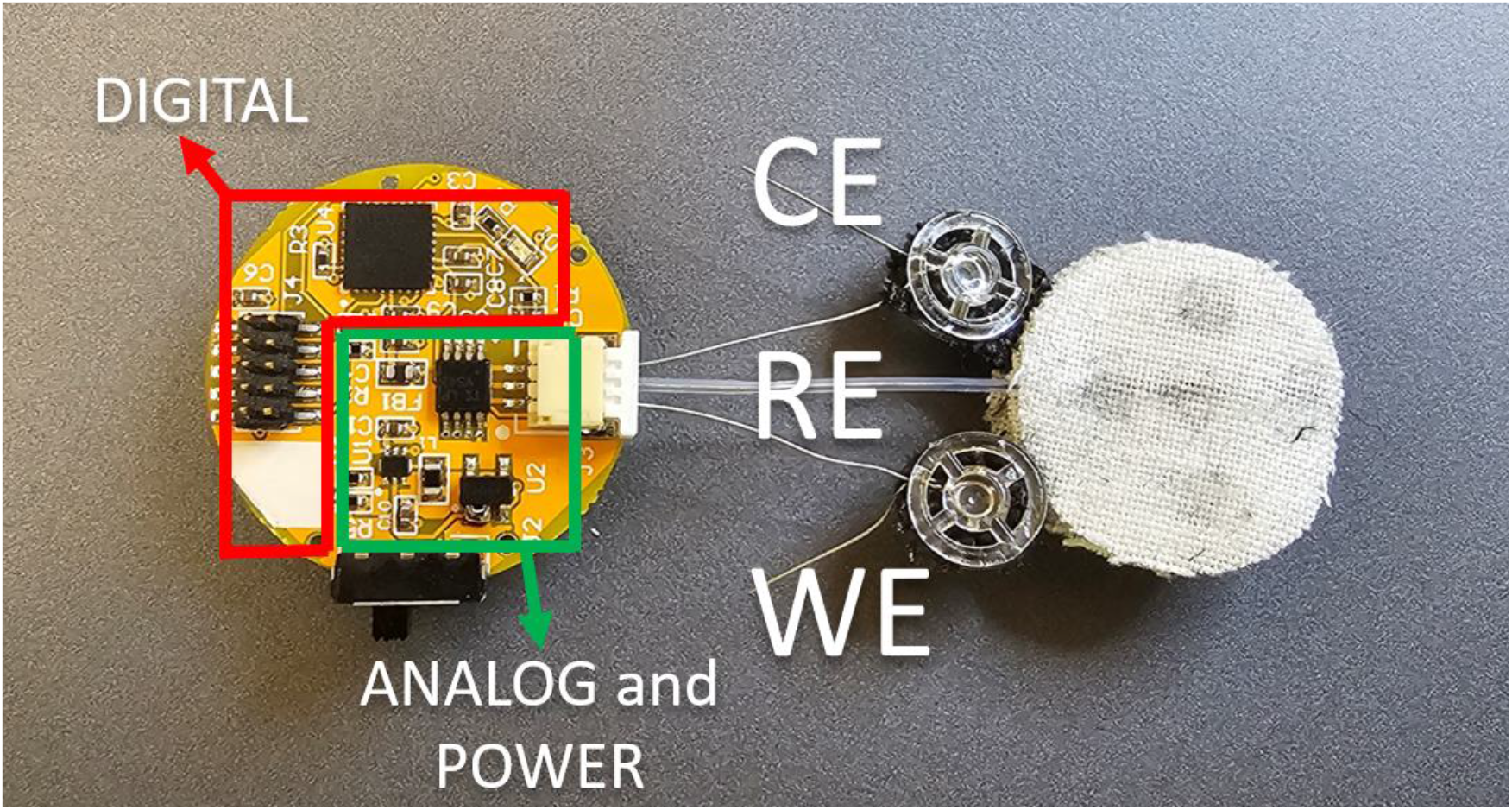
Photograph of an e-bandage connected to the PMWP. WE, RE, and CE correspond to the working electrode, reference electrode, and counter electrode of the e-bandage, respectively.

## Benchmark performance of the PMWP against a commercial potentiostat

### *Abiotic tests—*cyclic *voltammetry*

The performance of the PMWP was compared to that of a commercial potentiostat (Interface 1000E, Gamry Instruments, Warminster, PA, USA). The experimental setup involved a ferricyanide/ferrocyanide redox couple comprising 10 mM of potassium ferricyanide (K_3_Fe(CN)_6_) and 10 mM of potassium ferrocyanide (K_4_Fe(CN)_6_) in 100 mM of potassium chloride (KCl). An electrochemical reactor with a 30 mL working volume, featuring a glassy-carbon WE, a graphite CE, and a saturated Ag/AgCl RE was used. For benchmarking purposes, cyclic voltammetry was employed with scan rates of 5, 10, and 20 mV/s. Cyclic voltammograms (CVs) were generated with an initial potential of -0.7 V_Ag/AgCl_, using a 5-mV step size for each scan step. The upper boundary of the scan was set at +0.7 V_Ag/AgCl_. After reaching the maximum positive voltage, the scan returned to the initial voltage with set steps. The electrochemical parameters from the CVs included the anodic peak potential (*E*_*ρ,a*_), cathodic peak potential (*E*_*ρ,c*_), peak potential differences (*ΔE*), formal reduction potential (*E*°′), and anodic/cathodic peak current ratio (*i*_*ρ,a*_/*i*_*ρ,c*_). Calculation of these electrochemical parameters followed procedures outlined by Beyenal and Babauta (Beyenal and Babauta, 2015).

### Chronoamperometry and verification of H_2_O_2_ and HOCl generation

Custom-made microelectrodes were used to verify H_2_O_2_ or HOCl generation. Microelectrodes were constructed and calibrated following previously published protocols (Kiamco et al., 2019; Sournia-Saquet et al., 1999). Briefly, an e-bandage was placed onto a Petri dish in such a way that the WE faced upwards. Hydrogel was used to saturate the e-bandage, and a microelectrode was placed close to the WE surface. The WE was polarized to generate H_2_O_2_ or HOCl. The location of the microelectrode near the e-bandage surface was monitored using a stereomicroscope (Leica M80). The position of the microelectrode tip and its movement were controlled using a computer-linked stepper motor (Hysik Instrumente, PI M-230.10s, part no. M23010SX). The system was operated using custom LabVIEW script (National Instruments, Austin, TX, USA) software. To measure the H_2_O_2_ or HOCl concentration profile, 10-µm intervals were used. To measure H_2_O_2_ or HOCl concentration changes over time, the microelectrode was placed ∼50 µm from the WE surface. The e-bandage was operated by the PMWP and a Gamry Potentiostat Reference 600 (Gamry Instruments) while H_2_O_2_ or HOCl microelectrodes were operated by a Gamry Potentiostat Series G300 (Gamry Instruments).

### Testing broad-spectrum biocidal activity of PMWP

e-Bandages were built and tested following published procedures.(Mohamed et al., 2021, n.d.) e-Bandage biocidal efficacy was tested *in vitro* against 24-hour biofilms formed by clinical isolates of four prevalent chronic wound bacterial pathogens with concerning rates of antibiotic resistance,(“Prioritization of pathogens to guide discovery, research and development of new antibiotics for drug-resistant bacterial infections, including tuberculosis,” n.d.; Puca et al., 2021; Wolcott et al., 2016, p. 2) methicillin-resistant *Staphylococcus aureus* (MRSA), *Pseudomonas aeruginosa, Acinetobacter baumannii*, and *Enterococcus faecium*, and one fungal pathogen of emerging concern, *Candida auris*.(Lyman et al., 2023) (**Table 1**). Biofilms were grown on 13 mm polycarbonate membranes atop tryptic soy agar plates (TSA, Fisher Scientific, DF0370-17-3). Briefly, a single colony from an overnight agar streak plate was placed in two mL tryptic soy broth (TSB) within a 15 mL polystyrene Falcon tube and incubated, with shaking (120 rpm), at 37°C until the culture reached 0.5 McFarland standard. UV-sterilized 13 mm polycarbonate membranes were then placed on the TSA plates and inoculated with 2.5 µL of the culture. The plates were incubated at 37 °C for 24 hours to grow biofilms.

**Table 1.** Microbial strains studied.

| Species | Strain | Strain Details |
| --- | --- | --- |
| *Staphylococcus aureus* | IDRL-6169 | Prosthetic hip isolate; resistant to methicillin and mupirocin |
| *Enterococcus faecium* | IDRL-11790 | Abscess isolate; resistant to vancomycin and penicillin |
| *Pseudomonas aeruginosa* | IDRL-11442 | Groin isolate; resistant to piperacillin/tazobactam, cefepime, ceftazidime, meropenem, aztreonam, ciprofloxacin, and levofloxacin (colistin-susceptible) |
| *Acinetobacter baumannii* | IDRL-11626 | Wound isolate; resistant to amikacin, ampicillin, cefepime, ceftazidime, ciprofloxacin, and tobramycin |
| *Candida auris* | IDRL-12834 | Blood isolate; resistant to amphotericin B |

e-Bandages were polarized with the PMWP, to produce HOCl or H_2_O_2_ for desired portions of a 24-hour total treatment time. Treatment groups consisted of 1) 3 hours of HOCl production followed by 3 hours of non-polarization (no biocide production) and 18 hours of H_2_O_2_ production, 2) 3 hours of HOCl production followed by 21 hours of non-polarization, 3) 24 hours of HOCl production, 4) 24 hours of H_2_O_2_ production, 5) 6 hours of non-polarization followed by 18 hours of H_2_O_2_ production, 6) 24 hours of non-polarized control e-bandage, 7) no e-bandage control. Group 1 (3 hours HOCl, 3 hours non-polarized, 18 hours H_2_O_2_) was chosen to represent a theoretical scenario HOCl is produced for 3 hours, followed by 3 hours of HOCl dissipation (HOCl otherwise reacts with H_2_O_2_),(Anoy et al., 2022) and 18 hours of H_2_O_2_ delivery (to deliver a biocide and promote wound healing). After treatment, e-bandages were removed from TSA plates and colony forming units (CFUs) quantified via serial 1:10 dilution in PBS and drop plating on TSA plates.

## RESULTS AND DISCUSSION

### Electronic characterization of PMWP

Performance of an e-bandage controlled by a PMWP was evaluated by connecting the CE and RE and then placing a resistor (1-kΩ) between the two. This allowed PMWP performance to be tested under the most power-consuming conditions. **Fig. 3** shows the change in battery and cell voltages and sink current over time when the WE was polarized to 1.5 V_Ag/AgCl_. Considering that the values did not change during operation of up to 122 hours, only data for the first 1.5 hours are presented. The cell voltage remained constant while the battery voltage gradually dropped to 2.87 V. The sunk current gradually increased, reaching a plateau of 2.12 mA. Based on these tests, the PMWP could effectively operate for up to 122 hours when powered by a 260-mAh 2032-coin battery.

**Fig. 3.**
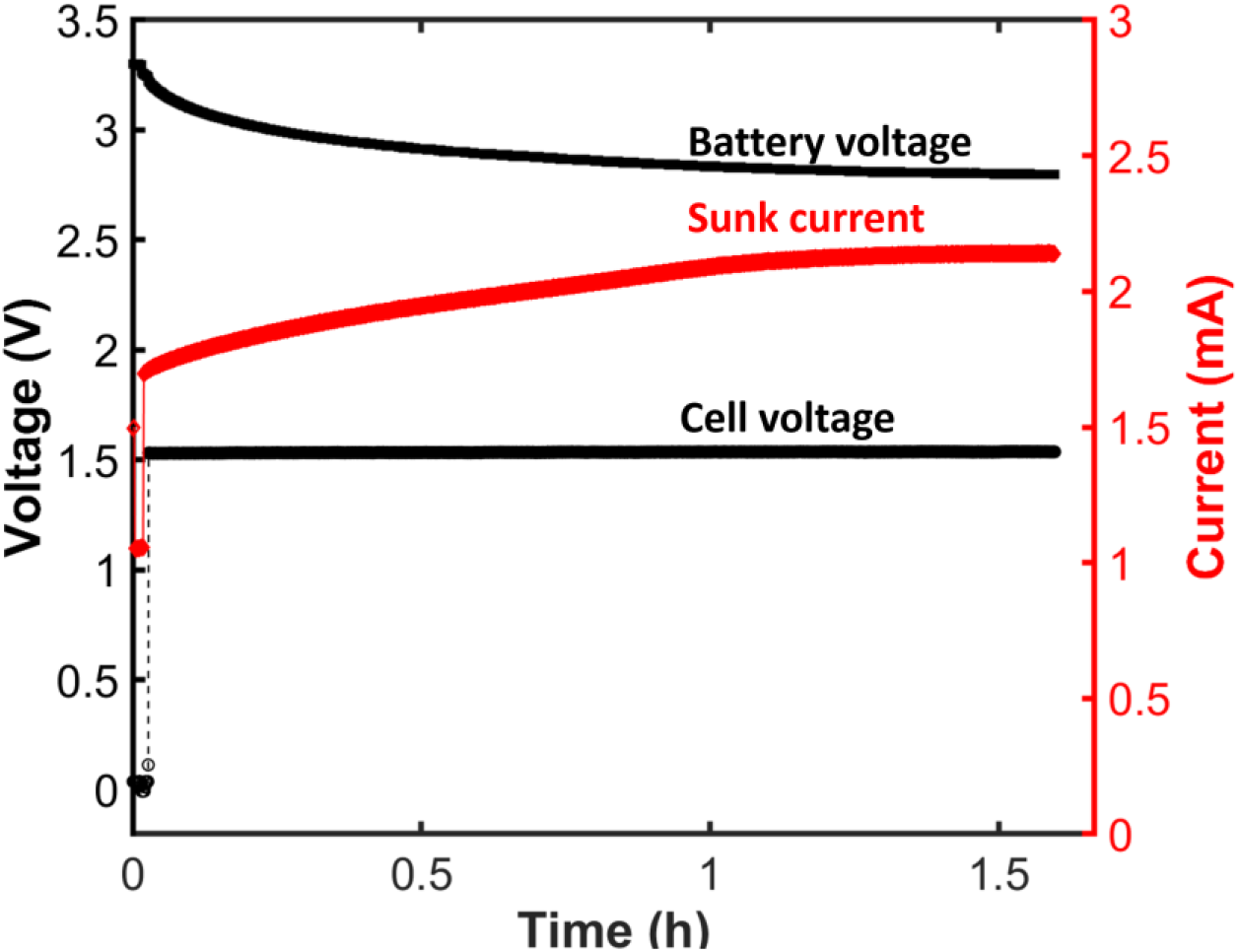
Change in battery and cell voltages and sunk current over time with the working electrode polarized to 1.5 V_Ag/AgCl_.

### Abiotic tests—cyclic voltammetry

Performance of the PMWP was tested in an abiotic electrochemical test, and its activity was compared to that of a commercial potentiostat (Gamry Instruments). **Fig. 4** shows cyclic voltammograms generated using the PMWP and a commercial potentiostat, as detailed in the Methods section, at scan rates of 5, 10, and 20 mV/s in ferricyanide/ferrocyanide redox couple. The anodic and cathodic peak currents rose with increasing scan rates. Additionally, the peak potential difference (difference between anodic and cathodic peak potentials) increased with a higher scan rate (**Fig. 4**). Finally, an increased scan rate led to an elevation of capacitive currents, which led to an increase in the hysteresis loop (**Fig. 4**). These phenomena are associated with the electrochemical cell, with the PMWP and the commercial potentiostat recording similar responses from the electrochemical cell. **Table 2** shows the anodic peak potential (*E*_*ρ,a*_), cathodic peak potential (*E*_*ρ,c*_), peak potential difference (*ΔE*), formal reduction potential (*E*°′), and anodic/cathodic peak current ratio (*i*_*ρ,a*_/*i*_*ρ,c*_) for numerical comparison. The two potentiostats showed identical responses under the same operational conditions. It should be noted that since PMWP can only run a single CV at a time, the first scan is shown.

**Table 2.**
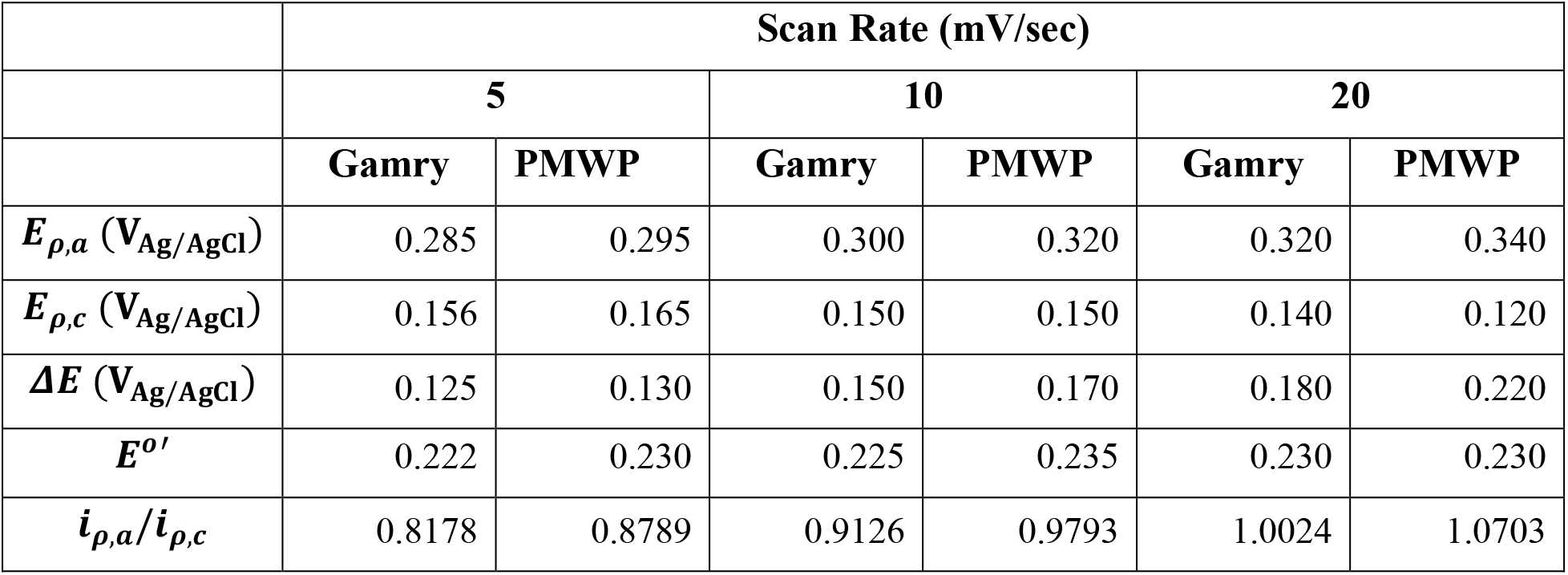
Parameters extracted from the scan rate analysis comparing CV performance of a commercial potentiostat (Gamry Instruments) and PMWP.

**Fig. 4.**
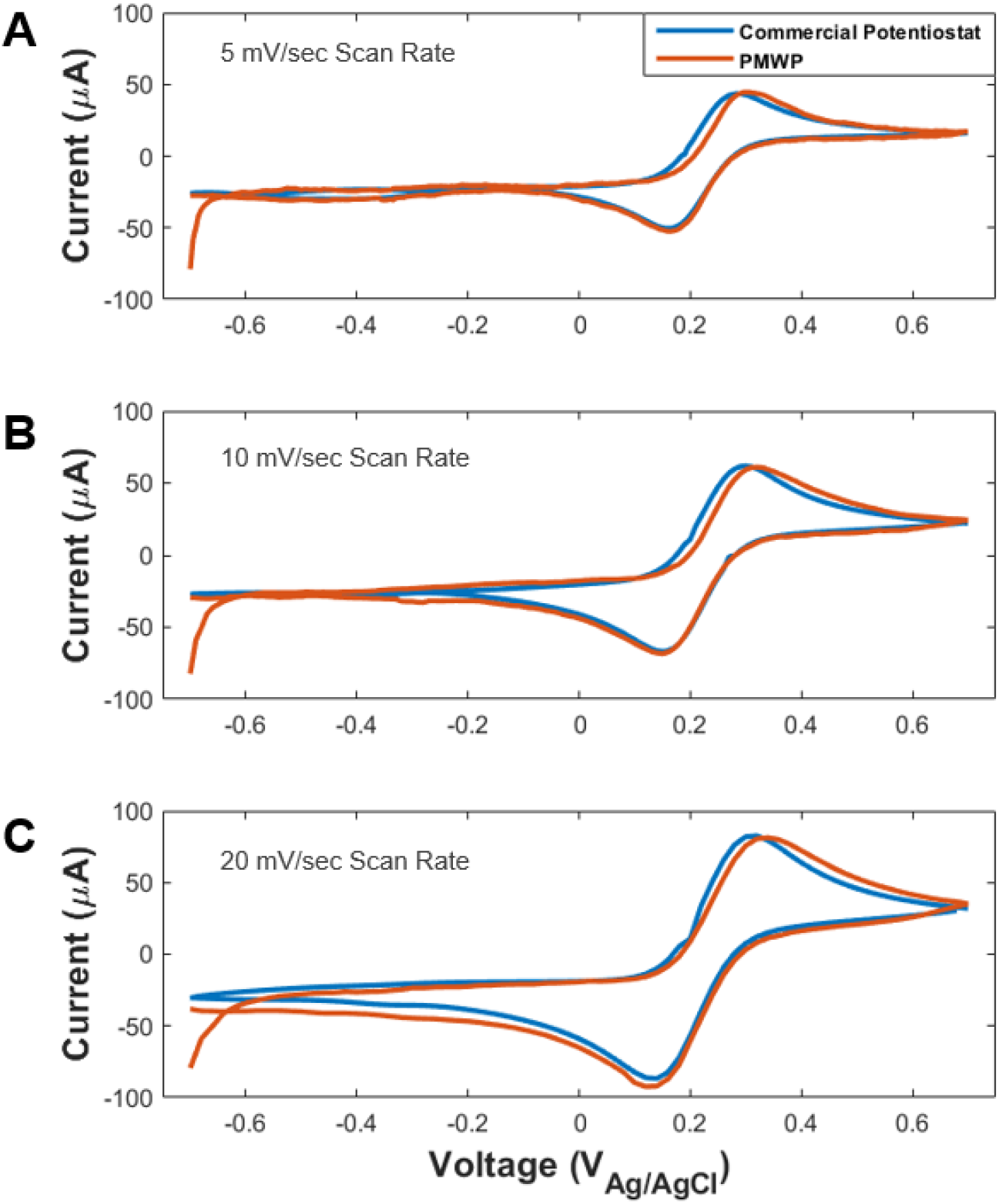
Comparison of PMWP and a commercial potentiostat in ferricyanide/ferrocyanide redox couple at (a, b, and c) scan rates of 5, 10, and 20 mV/s, respectively. PMWP and the commercial potentiostat (Gamry Instruments) recorded similar electrochemical behavior from the electrochemical cell.

### Verification of H_2_O_2_ and HOCl generation by PMWP-controlled e-bandages

Generation of H_2_O_2_ and HOCl by PMWP-controlled e-bandages was confirmed by measuring concentrations with microelectrodes situated near the electrode surface. In both cases, the PMWP was used to generate HOCl or H_2_O_2_ at the WE surface of an e-bandage. **Fig. 5A** and **5B** show changing HOCl and H_2_O_2_ concentrations measured ∼50 µm from the WE surface and the current consumed by the PMWP. Initially, concentrations were zero, increasing over time. HOCl concentrations reached a maximum of ∼900 µM and then decreased slowly because of the diffusion effect (**Fig. 5A**). A similar response was observed with H_2_O_2_ generation. The maximum H_2_O_2_ concentration was ∼360 µM (**Fig. 5B**). Both peak concentrations were observed after ∼15 minutes of PMWP operation. This was most likely due to the forward diffusion of HOCl or H_2_O_2_ to the bulk hydrogel. The depth profiles shown in **Fig. 5C** and **5D** indicate that the concentrations increase towards the WE surface. The fluxes of HOCl and H_2_O_2_ near the electrode surfaces were 2.96×10^-11^ and 2.34×10^-11^ moles/cm^2^/s, respectively. The data shown in **Fig. 5** demonstrate that the PMWP effectively generates HOCl and H_2_O_2_.

**Fig. 5.**
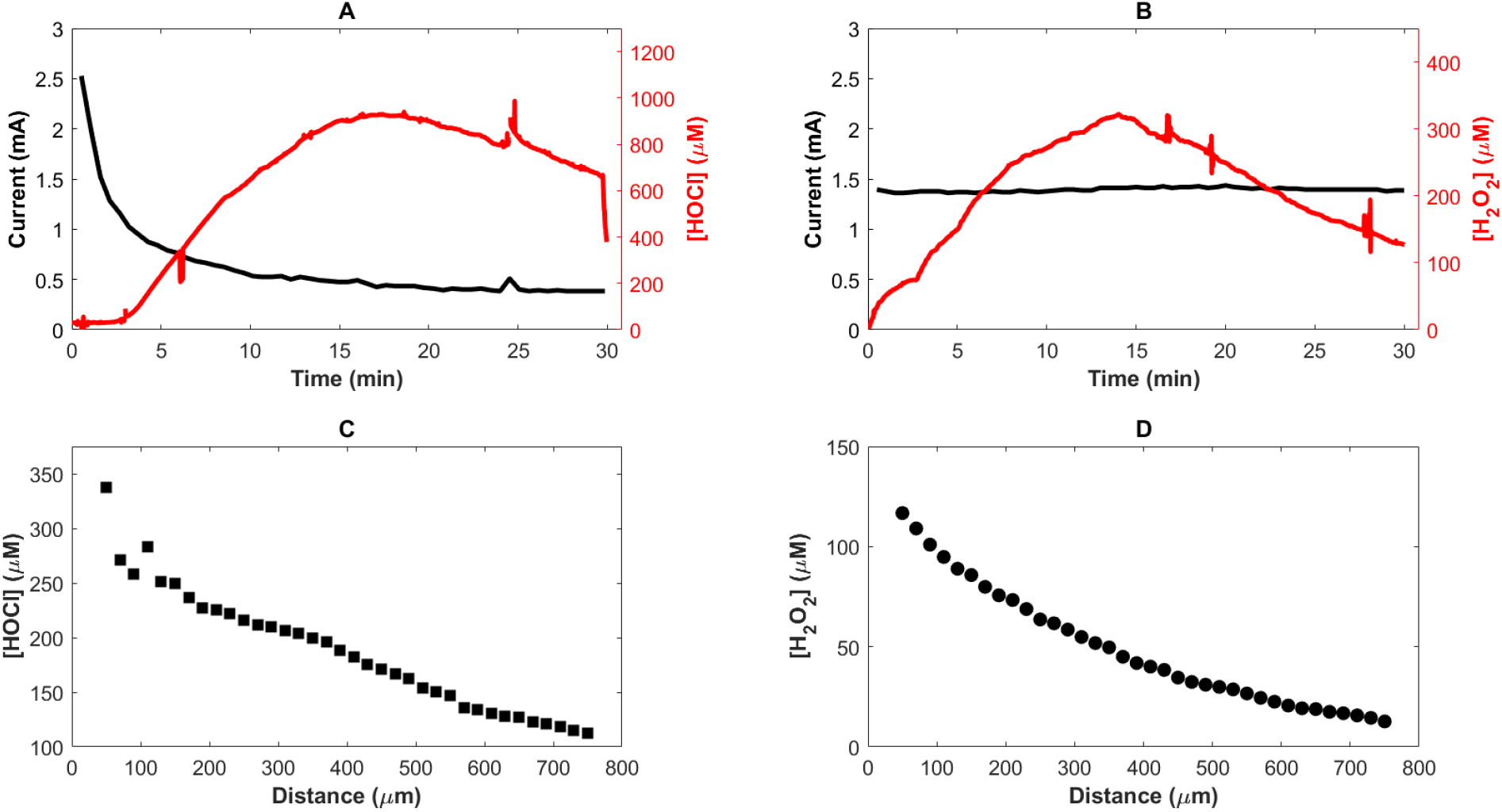
Verification of H_2_O_2_ and HOCl generation using PMWP. **A)** HOCl and **B)** H_2_O_2_ concentrations measured 50 µm from the working electrode (WE) surface and currents consumed during electrochemical reactions. The e-bandage was polarized at 1.5 V_Ag/AgCl_ for HOCl and -0.6 V_Ag/AgCl_ for H_2_O_2_. Depth concentration profiles within the hydrogel on top of **C)** HOCl**-** and **D)** H_2_O_2_-generating e-bandages after 30 minutes of operation. The zero-µm distance indicates the WE surface, and a positive distance value indicates distance from the WE surface.

### Biocidal activity of PMWP

The biocidal effectiveness of PMWP-controlled e-bandages against MRSA biofilms was tested in comparison to that of e-bandages powered by a commercial potentiostat (Gamry Instruments) (results not shown). Various treatment conditions were examined, and various durations of HOCl and H_2_O_2_ delivery were employed over six hours of total treatment time. Across all treatment conditions, there were no statistically significant differences between PMWP and the commercial potentiostat.

With the comparable biocidal activity of e-bandages controlled by PMWP and that of e-bandages controlled by a commercial potentiostat confirmed, the effectiveness of PMWP-controlled e-bandages against a range of clinically relevant wound pathogens was tested (**Fig. 6**). Clinical isolates with challenging antimicrobial resistance profiles were selected to demonstrate applicability of the e-bandage technology as a method to circumvent resistance. These pathogens included two Gram-positive (MRSA IDRL-6169 and *E. faecium* IDRL-11790) and two Gram-negative (*P. aeruginosa* IDRL-11442 and *A. baumannii* IDRL-11626) bacterial isolates recognized as prevalent bacterial pathogens with concerning rates of antibiotic resistance (“Prioritization of pathogens to guide discovery, research and development of new antibiotics for drug-resistant bacterial infections, including tuberculosis,” n.d.; Puca et al., 2021; Wolcott et al., 2016) and one fungal isolate, *C. auris*, that the World Health Organization (WHO) has identified as a pathogen of concern (Lyman et al., 2023). Twenty-four-hour membrane biofilms were treated with e-bandages polarized by PMWPs for 24 hours. Seven treatment groups were tested: 1) 3 hours of HOCl production, followed by 3 hours of non-polarization and then 18 hours of H_2_O_2_ production; 2) 3 hours of HOCl production followed by 21 hours of non-polarization; 3) 24 hours of HOCl production; 4) 24 hours of H_2_O_2_ production; 5) 6 hours of non-polarization followed by 18 hours of H_2_O_2_ production; 6) 24 hours of non-polarized e-bandage (control); and 7) no e-bandage (control). Groups 2-7 were compared to group 1 (the experimental group) to determine the efficacy of the theoretical idealized regimen. For all isolates tested, 24 hours of HOCl production resulted in elimination of the microbial population. In agreement with published results (Kletzer et al., 2023b; Mohamed et al., 2023), HOCl exhibited greater biocidal activity than H_2_O_2_ against all isolates. Interestingly, while HOCl exhibited biocidal efficacy against all study isolates, the ability of H_2_O_2_ to kill the biofilm pathogens varied, possibly because of differences in peroxidase activity between species and strains. For example, *E. faecium* showed a poor response to H_2_O_2_ alone and has been noted in the literature for its high peroxidase activity, especially NADH peroxidase, allowing it to reduce H_2_O_2_ to water (La Carbona et al., 2007). *P. aeruginosa* was similarly little affected by H_2_O_2_ and is known to possess an array of peroxidases and catalases (Sivaloganathan and Brynildsen, 2021).

**Fig. 6.**
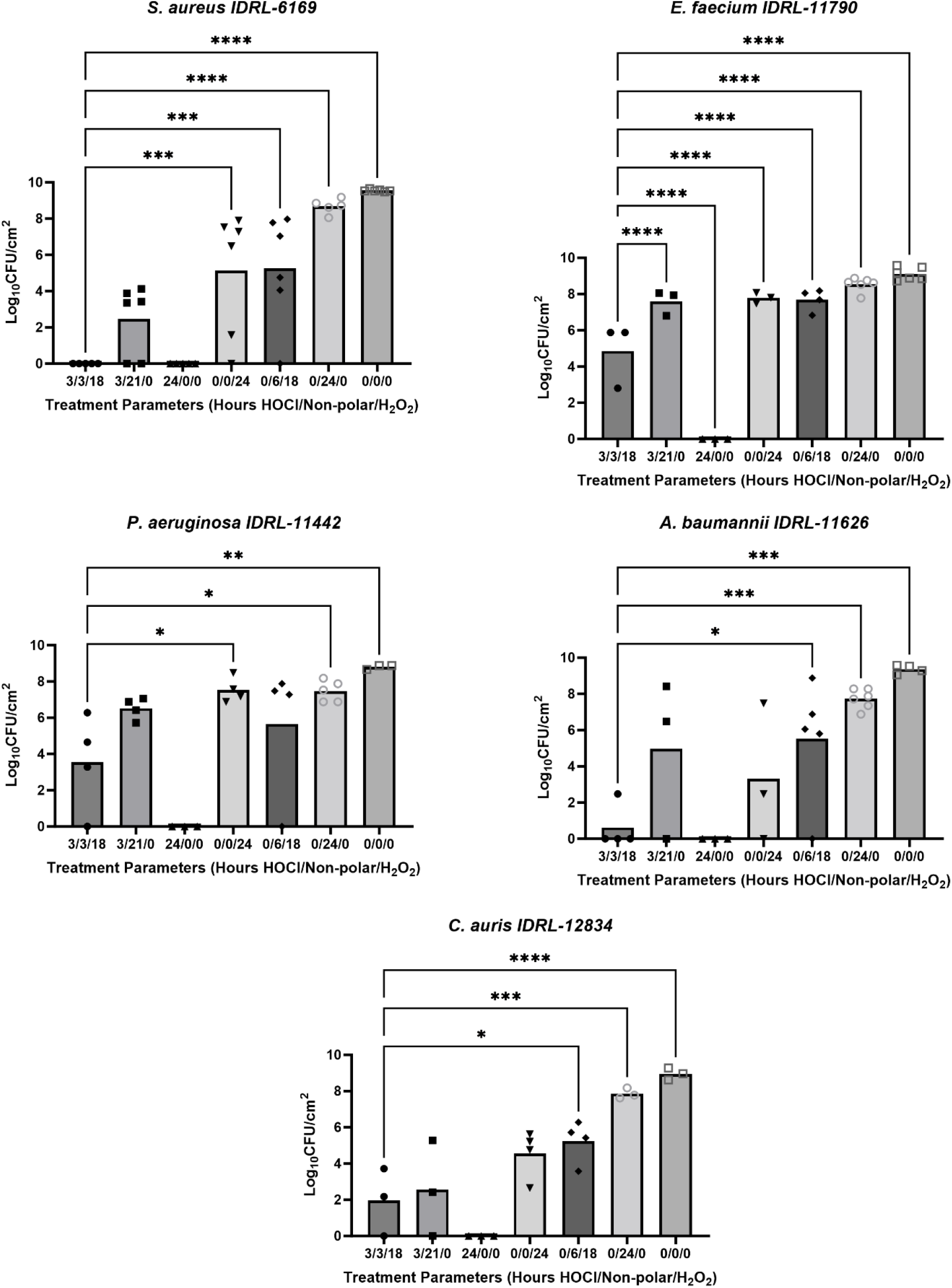
Efficacy of various PMWP-powered e-bandage treatment regimens with HOCl and H_2_O_2_ against bacterial and fungal pathogens. The e-Bandages were programmed with six different HOCl and H_2_O_2_ generation regimens with periods of non-polarization (represented on the x-axes as hours HOCl production/hours non-polarization/hours H_2_O_2_ production): 1) 3 hours of HOCl production followed by 3 hours of non-polarization (no biocide production) and 18 hours of H_2_O_2_ production (**3/3/18**), 2) 3 hours of HOCl production followed by 21 hours of non-polarization (**3/21/0**), 3) 24 hours of HOCl production (**24/0/0**), 4) 24 hours of H_2_O_2_ production (**0/0/24**), 5) 6 hours of non-polarization followed by 18 hours of H_2_O_2_ production (**0/6/18**), 6) 24 hours of non-polarized control e-bandage (**0/24/0**), and 7) no e-bandage control **(0/0/0**). Biofilm cell counts in CFUs are shown as log CFU/cm^2^. Data are represented as individual data points and means (bars) of at least three independent biological replicates (n≥3). All groups were compared to the theoretical clinical regimen (3/3/18) with one-way Anova with Dunn’s test for multiple comparisons; ^*^p≤0.05, ^**^p≤0.01, ^***^p≤0.001, ^***^p≤0.0001.

For MRSA biofilms, all groups including an HOCl production phase had biofilm reductions of at least 4.26 log_10_ CFU/cm^-2^ vs. no-e-bandage controls and at least 3.68 log_10_ CFU/cm^-2^ vs. non-polarized controls. While the groups containing HOCl production did not significantly differ from each other, group 1 showed more reduction than all groups not delivering HOCl. For *E. faecium*, group 1 resulted in significant reductions compared to all groups except the one with 24 hours of HOCl production. For *P. aeruginosa*, group 1 had significant reduction compared to all groups except those with 3 hours of HOCl production and 21 hours of non-polarization, 24 hours of HOCl production, or 6 hours of non-polarization followed by 18 hours of H_2_O_2_ production. Interestingly, despite the lack of reduction compared to 18 hours of H_2_O_2_ production, a significant reduction in group 1 did occur compared to 24 hours of H_2_O_2_ production. We note that this is likely due to a high standard deviation in the group polarized for 18 hours for H_2_O_2_ delivery. For both *A. baumannii* and *C. auris*, group 1 had significant reductions compared to all groups except those with 3 hours of HOCl production and 21 hours of non-polarization, 24 hours of HOCl production, or 24 hours of H_2_O_2_ production. These results, though variable, indicate that the PMWP-powered e-bandage is capable of broad-spectrum biofilm treatment and that a theoretical clinical application of a short HOCl delivery period followed by a prolonged period of H_2_O_2_ generation shows promise for continued testing in an animal wound infection model.

### Outlook

The PMWP is powered by a 260-mAh capacity coin battery and weighs 4.6 grams, making it suitable for small animal experiments and potentially human use. The PMWP can be operated without the battery needing to be replaced for 122 hours, and it can record current and potential data which can be retrieved for later analysis. The e-bandage is designed in such a way that the electronic current flows from the PMWP (or a commercial potentiostat) while ionic current moves within the hydrogel. Such a system does not allow current to move into animal or human tissues, making the device suitable for clinical applications. While other groups have proposed electric approaches for wound healing and/or wound infection treatment, the system described here is unique. Song et al. developed a miniaturized wireless, battery-free bioresorbable electrotherapy system which was evaluated on a murine wound model; while wound closure was improved, neither prevention nor treatment of infection was evaluated (Song et al., 2023) and their flexible system did not control WE voltage to generate biocides selectively. Instead, it utilized predefined voltage levels. Shirzaei Sani et al. developed a wearable patch to monitor physiochemical parameters (pH, NH_4_^+^, glucose) in real time in addition to applying voltage differences between two electrodes in a system using an antimicrobial peptide in hydrogel; <0.5 log_10_ reductions in CFUs were reported in a rat wound infection model (Shirzaei Sani et al., 2023). Heald et al. used a two-electrode system with 6 V applied between electrodes by limiting current (0.6 mA) to treat dog and cat wounds (Heald et al., 2022); no mechanism of action was proposed, so advancing the system to humans would be difficult. In contrast to these systems, the PMWP is mechanistically precise, allowing programmable generation of H_2_O_2_ or HOCl by the WE, and our previous iterations of single biocide-generating e-bandages have shown clinically relevant treatment activity in a murine wound infection model (Fleming et al., 2023; Raval et al., 2023). The PMWP allows for antimicrobial activity to potentially be coupled with wound healing augmentation. Another advantage of the PMWP is that it can be programmed for multiple desired applications apart from HOCl and H_2_O_2_ generation, from production of other chemicals to cyclic voltammetry for electrochemically monitoring wound conditions (**Fig. 5**).

In summary, the PMWP-powered e-bandage is a promising non-antimicrobial approach to wound infection treatment and wound healing.

## CONCLUSIONS

In this work, a PMWP was designed to operate an H_2_O_2_- or HOCl-generating e-bandage for treating biofilm infections. Electronic performance of the PMWP was tested, electrochemical performance and HOCl/H_2_O_2_ generation were verified, and the activity of PMWP-powered e-bandage treatment against *in vitro* biofilms was demonstrated. The PMWP-powered e-bandage showed similar biocidal activities to an e-bandage powered by a commercial potentiostat and exhibited broad-spectrum activity against wound infection-associated pathogens that exhibit antimicrobial resistance. The pathogenic strains tested represent clinical isolates with challenging resistance profiles, indicating that the PMWP-powered e-bandages are effective against even the most difficult biofilm infections to eradicate. Notably, PMWP-powered e-bandages delivering short-term HOCl production, followed by a washout period and then prolonged H_2_O_2_ production, were effective against all isolates tested, reducing biofilm populations by between 4.3 and 9.5 log_10_ CFU/cm^-2^ compared to no-bandage control groups.

## Supporting information

SI for MC paper

## ACKNOWLEDGEMENTS

Research reported in this publication was supported by the National Institute of Allergy and Infectious Diseases of the National Institutes of Health under award number R01 AI091594. PFR was supported by National Institutes of Health T32GM145408. The content is solely the responsibility of the authors and does not necessarily represent the official views of the National Institutes of Health.

