## Supplementary material for "DUAL ACTION ELECTROCHEMICAL BANDAGE OPERATED by a PROGRAMMABLE MULTIMODAL WEARABLE POTENTIOSTAT": SI for MC paper

Ibrahim Bozyel (0000-0002-2841-7890) ^a^, Derek Fleming (0000-0002-0054-904X)^b~^, Won-Jun Kim (0000-0003-3948-7672) ^a~^, Peter F. Rosen (0000-0003-4541-7283) Suzanne Gelston (0000-0001-6853-8067)^a^, Dilara Ozdemir^a^, Suat U. Ay (0000-0001-7640-4253) ^c^, Robin Patel (0000-0001-6344-4141)^b,d^, Haluk Beyenal (0000-0003-3931-0244) ^a,*^

^a^The Gene and Linda Voiland School of Chemical Engineering and Bioengineering, Washington State University, Pullman, WA, USA

^b^ Division of Clinical Microbiology, Mayo Clinic, Rochester, Minnesota, USA

^c^Department of Electrical and Computer Engineering, Worcester Polytechnic Institute, Worcester, MA, USA

^d^Division of Infectious Diseases, Mayo Clinic, Rochester, Minnesota, USA

^~^ These authors contributed equally

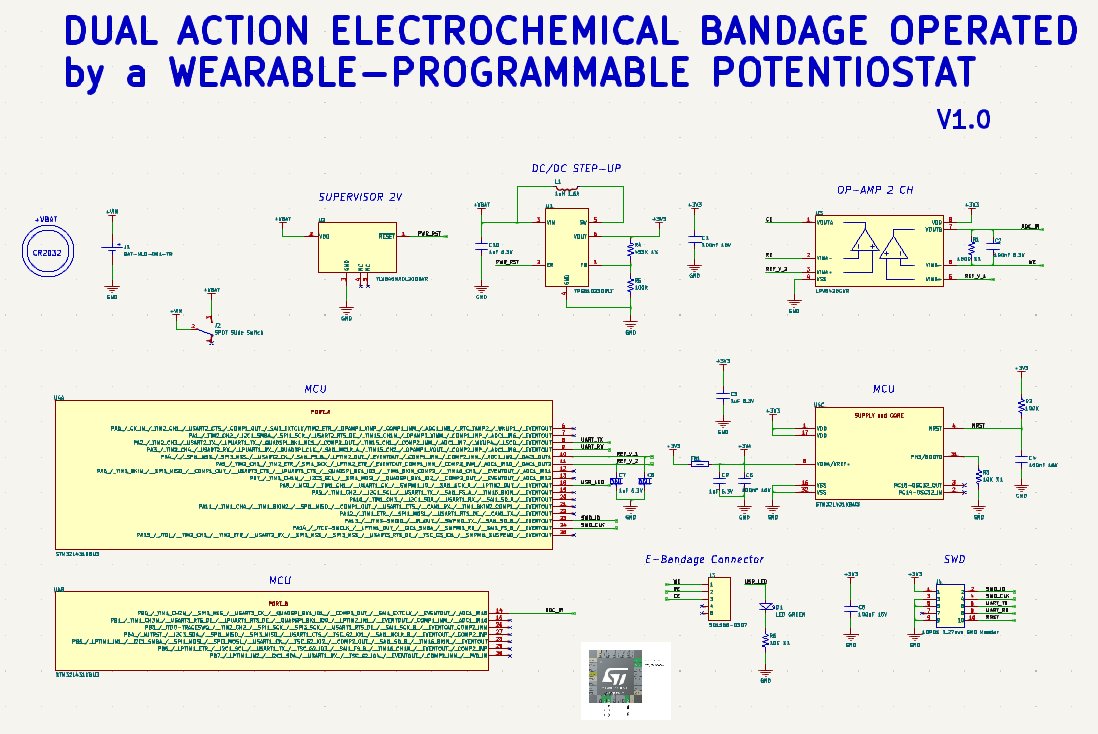

**Fig. S1.** Schematic design of PMWP. Designed using KiCAD.

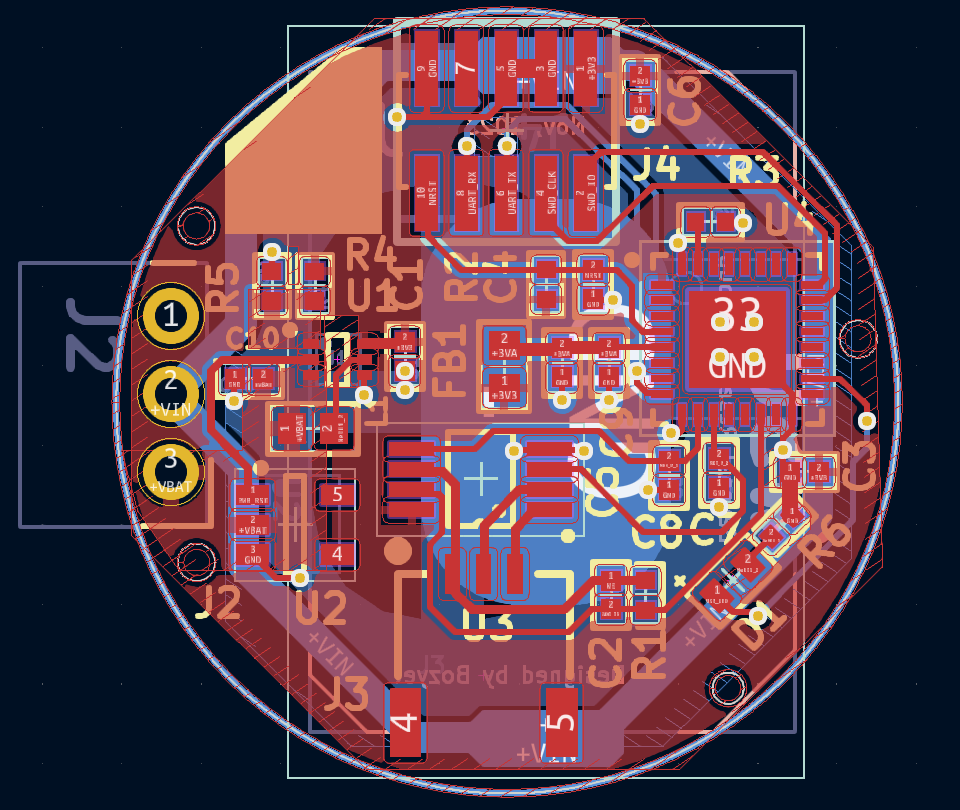

**Fig. S2.** Component placement and layout design of PMWP. Red indicates the top layer; blue indicates the bottom layer. Designed using KiCAD.
